# Fast and flexible bacterial genomic epidemiology with PopPUNK

**DOI:** 10.1101/360917

**Authors:** John A. Lees, Simon R. Harris, Gerry Tonkin-Hill, Rebecca A. Gladstone, Stephanie W. Lo, Jeffrey N. Weiser, Jukka Corander, Stephen D. Bentley, Nicholas J. Croucher

## Abstract

The routine use of genomics for disease surveillance provides the opportunity for high-resolution bacterial epidemiology.

However, current whole-genome clustering and multi-locus typing approaches do not fully exploit core and accessory genomic variation, and cannot both automatically identify, and subsequently expand, clusters of significantly-similar isolates in large datasets and across species.

Here we describe PopPUNK (Population Partitioning Using Nucleotide K-mers; https://poppunk.readthedocs.io/en/latest/). software implementing scalable and expandable annotation- and alignment-free methods for population analysis and clustering.

Variable-length *k*-mer comparisons are used to distinguish isolates’ divergence in shared sequence and gene content, which we demonstrate to be accurate over multiple orders of magnitude using both simulated data and real datasets from ten taxonomically-widespread species. Connections between closely-related isolates of the same strain are robustly identified, despite variation in the discontinuous pairwise distance distributions that reflects species’ diverse evolutionary patterns. PopPUNK can process 10^3^-10^4^ genomes as single batch, with minimal memory use and runtimes up to 200-fold faster than existing methods. Clusters of strains remain consistent as new batches of genomes are added, which is achieved without needing to re-analyse all genomes de novo.

This facilitates real-time surveillance with stable cluster naming and allows for outbreak detection using hundreds of genomes in minutes. Interactive visualisation and online publication is streamlined through automatic output of results to multiple platforms.

PopPUNK has been designed as a flexible platform that addresses important issues with currently used whole-genome clustering and typing methods, and has potential uses across bacterial genetics and public health research.

## Introduction

Determining whether a set of pathogen isolates are significantly more genetically similar than randomly-selected representatives from the circulating population is critical in identifying transmission pairs, localised outbreaks or global patterns of dissemination (Croucher and Didelot 2015). For phenotypically diverse bacterial pathogens, categorising sets of similar isolates is particularly valuable, as such clusters are often strongly associated with variation in clinically-relevant traits, including host range (Weinert et al. 2015; Reuter et al. 2014; Willems et al. 2012), virulence (Alikhan et al. 2018; Reuter et al. 2014; Weinert et al. 2015), propensity to cause nosocomial outbreaks (Willems et al. 2012; Aanensen et al. 2016) and antimicrobial resistance profile (Aanensen et al. 2016; Kallonen et al. 2017). Following the trends in these clusters over time provides critical information on population-level changes following the emergence of new strains (Kallonen et al. 2017), or resulting from interventions such as vaccine introduction (Croucher et al. 2013). Therefore the clustering of isolates into strains, and ascertaining the higher-resolution structure within these groupings, is a critical challenge in pathogen population genetics.

The earliest bacterial typing approaches used phenotypic assays, such as antibody binding (serotyping), phage infection (phage typing) or metabolic properties (biotyping) (Barker and Old 1989). However, these categorisation methods were typically genus- or species-specific, and limited in resolution by the number of detectable outcomes. Later epidemiological studies instead favoured methods that could be generalised across multiple bacteria and which were designed to assay selectively neutral variation across multiple loci, thereby avoiding the risk of being confounded by a single recombination or selective sweeps. These typically employed gel electrophoresis to distinguish between alleles within conspecific genotypes, either based on protein charge (multilocus enzyme electrophoresis, MLEE), tandem repeat length (multilocus variable number tandem repeat analysis, MLVA), or variation in restriction site distributions (pulsed-field gel electrophoresis, PFGE). However, despite the the standardisation of such approaches by PulseNet International (Swaminathan et al. 2001, 2006), comparison of gels between laboratories remains difficult.

Sequence data has the advantage of being more portable owing to its ease of standardisation and digitisation. Consequently, multilocus sequence typing (MLST) emerged as the gold standard epidemiological typing for bacteria in the late 1990s (Maiden et al. 1998). MLST labels are defined by any set of unique sequences at several short fragments of unlinked housekeeping genes; treating any change regardless of number of polymorphisms as the same provides robust clustering in the presence of recombination (Feil et al. 2004; Turner et al. 2007). Continually-updated online MLST databases have facilitated rapid comparisons between global isolate sets collected over decades (Jolley et al. 2017; Aanensen and Spratt 2005). Furthermore, as each MLST label is made up from several separate allele labels from each gene in the scheme, minimum-spanning trees can be generated using eBURST (Feil et al. 2004). This allowed sequence types to be grouped together into clonal complexes. However, the fixed resolution of MLST has two problems: firstly, pathogens such as *Mycobacterium tuberculosis* and *Salmonella enterica* serovar Typhi exhibit so little variation that MLST often struggles to distinguish conspecific isolates (Achtman 2012). Secondly, at the other extreme, some pathogens recombine sufficiently frequently that the limited resolution of MLST causes infrequent spurious links between unrelated groups of isolates, resulting in ‘straggly’ clonal complexes encompassing highly divergent bacteria (Turner et al. 2007; Willems et al. 2012).

Whole genome sequence data provides an opportunity to greatly improve the precision and resolution of bacterial typing. Core genome MLST (cgMLST) schemes have been designed as an extension to MLST by applying the same approach across the core genome, and have demonstrated their value at scales ranging from genus-wide taxonomy to investigation of nosocomial outbreaks in *Neisseria* spp. (Bratcher et al. 2014), *Listeria monocytogenes* (Ruppitsch et al. 2015), *Enterococcus faecium (De Been et al. 2015), Escherichia coli, Pseudomonas aeruginosa, Klebsiella pneumoniae* and *Staphylococcus aureus* (Mellmann et al. 2016). The advantage of these schemes is the speed and ease of assigning indices to alleles combined with the increased resolution of using larger proportions of the core genome. However, all such analysis are contingent upon, and limited to, the coding sequences identified in the original scheme: in the species-specific schemes listed above, this varied between 41% and 84% of the genes in a typical genome. This can only be achieved if there has been complete assembly of all these loci in the query genome. Further resolution is lost if these data are treated as a set of allele identifiers, rather than nucleotide sequences, as this obscures the level of similarity between non-identical alleles. Nevertheless, cgMLST is highly sensitive and can uncover deeper relationships between strains. This is a trade-off with specificity, meaning a minimum-spanning tree constructed using sequence types is fully connected, losing the simple and intuitive splitting of the population into clonal complexes (Feil et al. 2004). Some cgMLST studies have therefore resorted to using complexes from MLST (Maiden and Harrison 2016; Alikhan et al. 2018), with which they are highly consistent.

Subdivision of genetically diverse populations is critical for phylogeny-based analyses, such as phylodynamic studies (Weinert et al. 2015; Croucher et al. 2013; Kallonen et al. 2017; Kremer et al. 2017) or recombination identification (Croucher et al. 2013; Weinert et al. 2015). The identification of clusters from genomic data can be performed by partitioning a phylogeny (Prosperi et al. 2011), but has more typically used complex population structure analysis models: initially STRUCTURE (Pritchard et al. 2000) or BAPS (Corander et al. 2008), and more recently hierBAPS (Cheng et al. 2013). Comparisons between the clusters generated by these algorithms and MLST clonal complexes again found high levels of overlap (Alikhan et al. 2018; Aanensen et al. 2016; Kallonen et al. 2017; Croucher et al. 2013), indicating these groupings correspond to natural bacterial populations. These methods fully exploit the available polymorphic sites extracted from core genome alignments, though these can be computationally challenging to generate for large or diverse population samples. There are two additional difficulties encountered when applying these sophisticated analytical approaches to pathogen surveillance. The first is that the identifiability of clusters depends on their frequency in the population, with rarer strains often either grouped together or treated as admixtures of more common genotypes (Grad et al. 2016; Willemse et al. 2016; Croucher et al. 2013), making changes in prevalence difficult to track prospectively. Secondly, these methods have to be rerun from scratch every time the alignment changes, which is further complicated due to their large computational burden, meaning they lack the simple extendability of MLST-type schemes.

Given the issues with core genome-dependent approaches, recent work has sought to improve the resolution of expandable typing schemes by exploiting the variation in accessory loci. Differences in gene content underlie much phenotypic variation between bacteria, and these are correlated with core genome divergence in multiple species (Croucher et al. 2014b; Zhou et al. 2014; Holt et al. 2015; Aanensen et al. 2016), motivating the use of this information in epidemiological typing. Hence whole genome MLST (wgMLST) schemes have been developed to incorporate accessory genes, by adding an allele identifier which corresponds to its absence, effectively weighting the acquisition of a single gene the same as individual SNPs (Liu et al. 2016). However, as variation in mobile elements can import many genes over short timescales (Abudahab et al. 2017; Croucher et al. 2016) and potentially be confounding for resolving outbreaks (Zhou et al. 2013), it can be preferable to keep such variation distinct from that in the core genome. Additionally, unlike the well-defined and conserved loci comprising cgMLST schemes, many wgMLST loci are more difficult to align and define, and can be complicated by the difficulty of resolving orthologous and paralogous genes (Zhang et al. 2015). Additionally, the extensive gene content variation of many species means a fixed scheme designed only using an initial sample will miss many accessory genes, including any newly-emerged loci that enter the population through horizontal gene transfer (Alikhan et al. 2018), the detection of which represents a critical aspect of pathogen surveillance. Alternatively, wgMLST schemes can be continually updated through searching each new isolate for previously-unseen loci, but this necessitates every genome being compared to a continually-expanding set of typically rare loci, accompanied by annotation and extraction of any new genes to add to the scheme, which is both computationally-intensive and algorithmically complicated.

Overall, typing based approaches such as MLST and its extensions suffer either from sub-optimal resolution or problems identifying biologically-meaningful clusters, whereas model-based population structure analysis is not suited to surveillance due to its computational burden and difficulty to extend to new isolates. To solve these methodological difficulties in a single approach, we have developed PopPUNK (Population Partitioning Using Nucleotide K-mers). By using efficient MinHash based comparisons between pairs of genome assemblies with *k*-mers (strings of bases of length *k*) of different lengths, we can distinguish divergence in orthologous sequence and variation in gene content. These distances are automatically and rapidly clustered to define strains, which we define here as sets of isolates significantly similar in both their core and accessory genomes relative to the rest of the species, in a frequency-independent manner. We have tested PopPUNK on simulated data and real datasets from ten bacterial species, demonstrating that the software rapidly and accurately estimates genetic distances between isolates. These are used to define clusters within samples more quickly than existing methods. Using network-based approaches, PopPUNKs output can be easily queried and expanded, meaning it is able to fully exploit genomic data in a scalable manner appropriate for real-time surveillance of large, diverse bacterial populations.

## Results

### PopPUNK uses variable length *k*-mers to accurately resolve genetic divergence

We proposed the probability that a *k*-mer will match between a pair of sequences, *p*_match_, is the product of accessory and core mismatches. Specifically, *p*_accessory_ the probability it does not represent an accessory locus unique to one member of the pair, and *p*_core_, the probability it represents a shared core genome sequence that does not contain any mismatches. To calculate *p*_core_ and *p*_accessory_, comparisons were run using the MinHash algorithm (Broder 1997) as implemented in Mash (Ondov et al. 2016), which estimates the Jaccard similarity between reduced size *k*-mer ‘sketches’ of the two sequences. This is run for a selection of *k*-mer lengths between *k*_min_ and *k*_max_, the former being determined by the minimum sequence length needed to avoid frequent false positive matches given the size of the genomes being compared (see Methods), and the latter by limited by memory-efficient MinHash processing (PopPUNK uses 29 by default). By determining the probability of *k*-mer matching *p*_core_ over this *k*-mer size range, it is possible to estimate the density of single nucleotide polymorphisms (SNPs) distinguishing the pair across their shared core, defined as π (Nei and Li 1979). This is because longer *k*-mers are more likely to contain a SNP, as quantified by the probability of a *k*-mer perfectly matching between the pair:

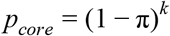

This approach assumes a random and uniform distribution of SNPs across the core, which is defined as those genomic regions in which nucleotide strings at least *k*_min_-long can be matched, representing statistically significant similarity between the pair. Loci in which there are no *k*_min_-long matches, resulting from either absence of the sequence in one member of the pair, or high sequence divergence of at least one SNP per *k*_min_ bases, are classified as belonging to the accessory genome; *k*_min_ thereby provides an intrinsic statistical distinction between the core and accessory regions in the pairwise comparison. Hence *p*_accessory_ can be regarded as the Jaccard similarity between a pair in terms of their shared core sequence, allowing the definition of the accessory divergence between sequences, a, as the corresponding Jaccard distance:

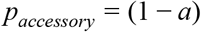

Unlike *p*_core_, *p*_accessory_ is independent of *k*, allowing both π and a to be jointly estimated from the assumption thatp_maŧch_ is the product of *p*_match_ and *p*_accessory_ (Fig 1):

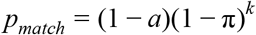

**Figure 1:**
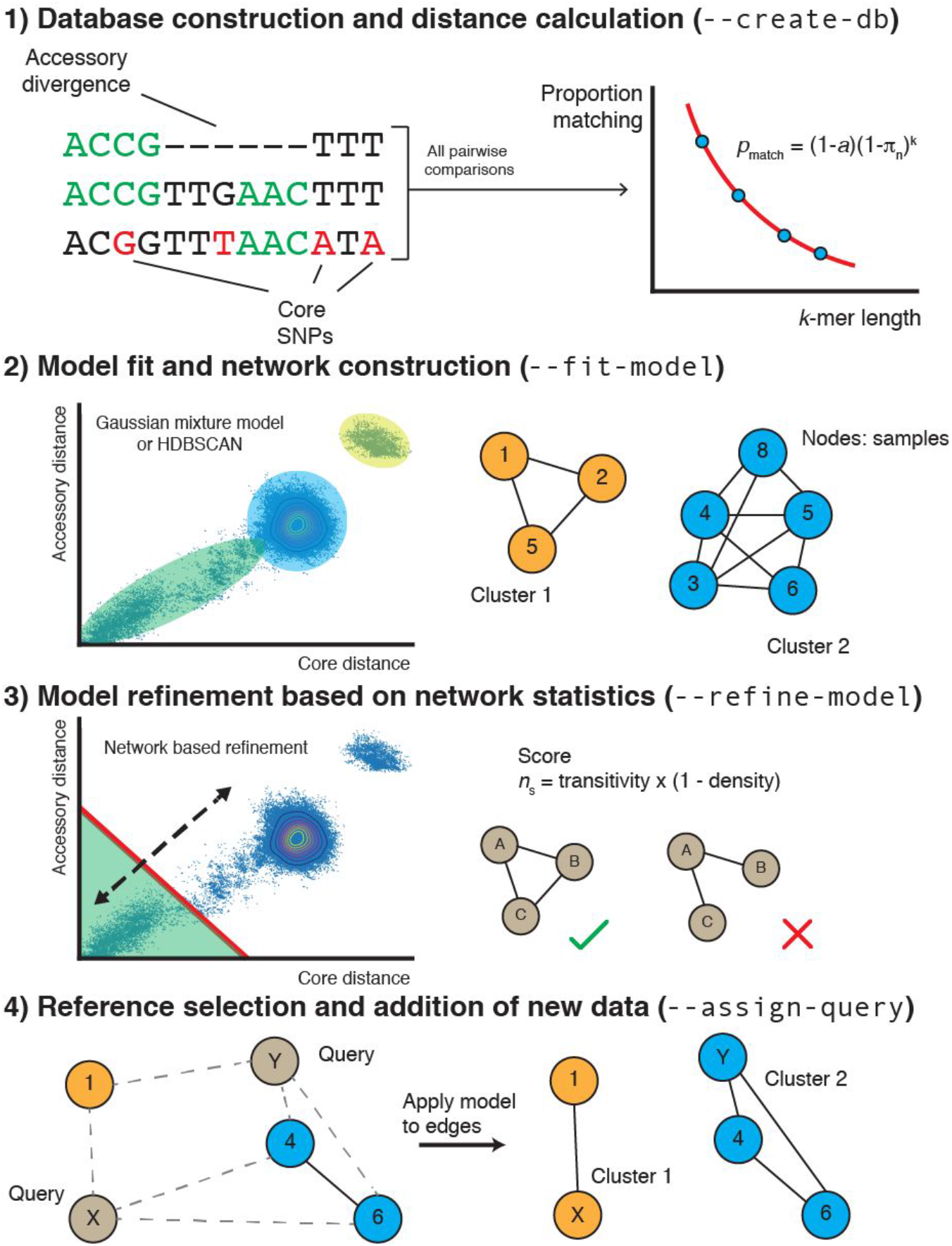
Summary of the PopPUNK algorithm. Step 1: For each pairwise comparison of sequences, the proportion of shared *k*-mers of different lengths is used to calculate a core and accessory distance. We use the fact that differences in gene content cause *k*-mers (examples highlighted in green) to mismatch irrespective of length, and whereas substitutions distinguishing orthologous sequences cause longer *k*-mers to mismatch more frequently than shorter *k-*mers. Step 2: The scatter plot of these core and accessory distances is clustered to identify the set of distances representing ‘within-stain’ comparisons between closely-related isolates. A network is then constructed from nodes, corresponding to isolates, linked by short genetic distances, corresponding to ‘within-strain’ comparisons. Connected components of this network define clusters. Step 3: The threshold defining within-strain links is then refined using a network score, n_s_, in order to generate a sparse but highly clustered network. Step 4: Finally, the network is pruned by taking one sample from each clique. The distances between new query sequences and references are calculated, and ‘within-strain’ distances used to add new edges. The clusters are then re-evaluated as in step 3, with the nomenclature being kept consistent with the original reference cluster names.

To test whether this approach was effective in differentiating core and accessory divergences, we performed forward-time simulations of bacterial populations diversifying through base substitutions, large insertions and deletions (indels) and recombination using BacMeta (Sipola et al. 2018). Those simulations in which sequences diverged only through base substitutions all correctly identified a < 5×10^−3^, whereas π increased according to the set substitution rate over multiple orders of magnitude, even in the presence of recombination (Fig 2A and S1). To test the accuracy with which a can be estimated, the substitution rate was fixed at 5×10^−6^ bp^−1^ generation^−1^, and indels of a fixed size of 250 bp occured at varying rates relative to base substitutions to emulate changes of gene content. The calculated a co-varied with the indel rate, without substantially affecting the inferred distribution of π (Fig 2B and S1). With the indel rate was fixed at 0.05, the distributions of both a and π converged towards a single mode as the rate of exchange through recombination was increased (Fig 2C), consistent with changes in the analogous core and accessory genetic distances observed in a study using a different framework to study the effects of sequence exchange (Marttinen et al. 2015). Hence PopPUNK’s use of variable length *k*-mers can resolve variation in the genome content and sequence to generate a pairwise distance distribution that accurately reflects the population-wide distribution of genetic diversity.

**Figure 2:**
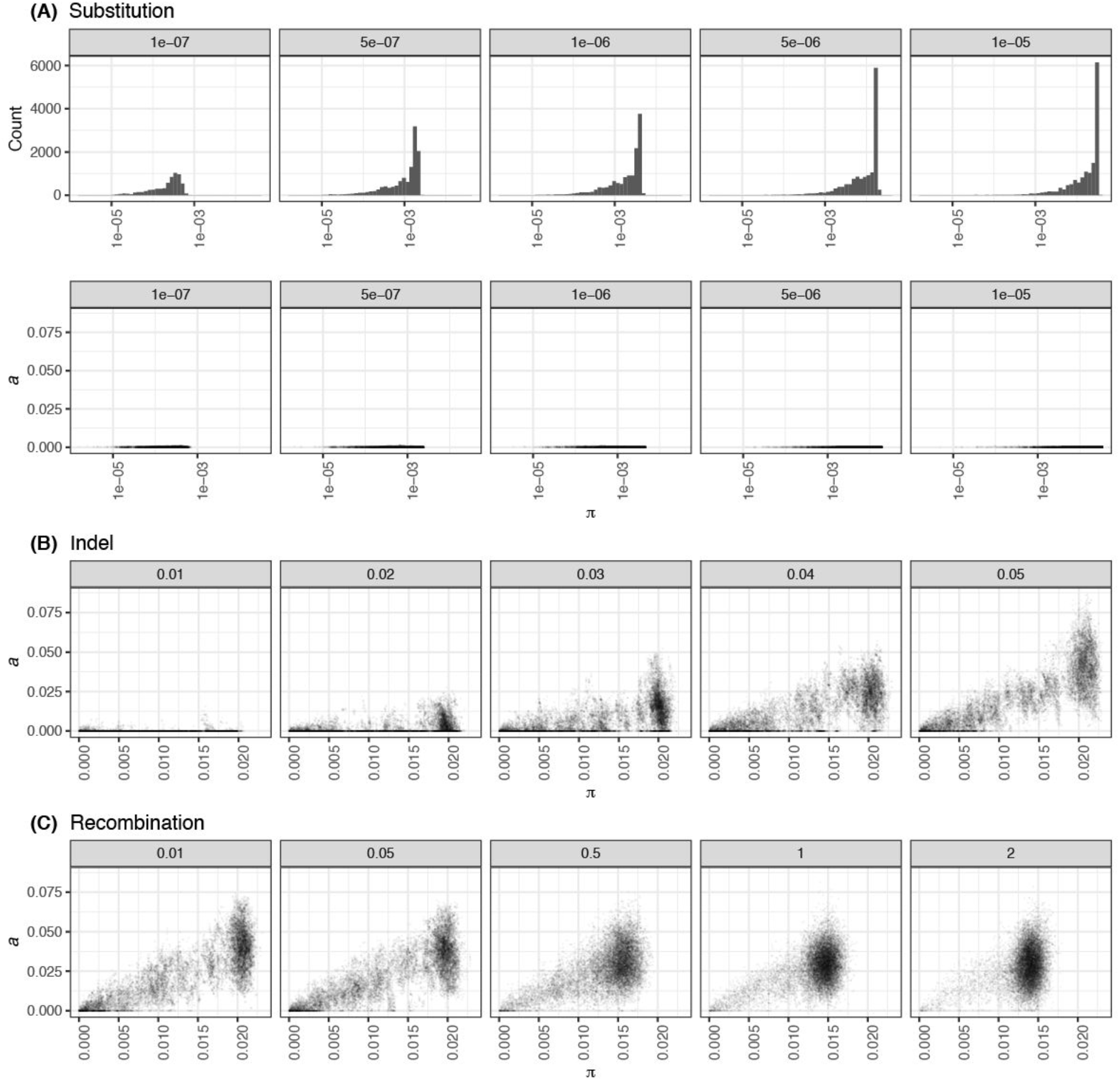
Detection of genetic diversity by PopPUNK in simulated populations. Each plot shows the deviation in gene sequence (π) and gene content (a) estimated by PopPUNK from a sample of 25 isolates from each of 50 simulations run with the same parameters. (A) Deviation through base substitution only. As the rate of base substitution (base^−1^ generation^−1^) was increased over two orders of magnitude, estimates of population-wide π increased accordingly, as shown by the distribution of pairwise core distances in the top row of histograms. The scatterplots beneath show a measurements remained below 5×10^−3^, demonstrating the specificity with which divergence was measured. (B) Deviation through insertions and deletions. To test the estimation of a in a clonally-evolving population, simulations included insertions and deletions of 250 bp segments occurring at a rate defined relative to the fixed substitution rate of 5×10^−6^ base^−1^ generation^−1^. Estimates of a increased proportionately with this rate, without affecting the observed range of π. (C) Effects of recombination on the distribution of genetic diversity. With the insertion and deletion rate fixed at 0.05 relative to the substitution rate of 5×10^−6^ base^−1^ generation^−1^, the rate of recombination relative to base substitutions was then varied. This resulted in a concentration of the estimated distances into a single mode, representing the changing population structure as frequent exchange between isolates homogenises the divergence between them in both gene sequence and content.

To determine the limits of resolution possible using PopPUNK, and therefore whether it could be used for surveillance of monomorphic pathogens or clonally-related outbreaks, distances were calculated between eight artificially-generated variants of a 2.2 Mb *Streptococcus pneumoniae* genome distinguished by three biallelic SNPs (Fig S2). This necessitates the use of a sufficiently large sketch size (10^5^), which determines the coverage of the genome in the *k-*mer representation. But with this increase even sequences distinguished by only a single SNP could be successfully resolved using a MinHash approach. This necessitates slightly longer runtimes, so we allow the user to select a larger sketch size than the default if there is a low level of divergence expected between isolates. Therefore given sufficiently high-quality data, PopPUNK can accurately resolve bacterial sequences at multiple scales of genetic divergence.

### PopPUNK identifies divergence between bacterial genomes across multiple species

To test whether PopPUNK could also produce accurate estimates of a and π when applied to real high-throughput sequencing data, the software was next applied to recent population genomics studies from ten diverse bacterial species. These were chosen to have varied ecologies, including enteric bacteria (*Escherichia coii* and *Salmonella enterica*), Gram negative respiratory pathogens (*Haemophilus influenzae* and *Neisseria meningitidis*), streptococci (S. *pneumoniae* and *Streptococcus pyogenes*), other Firmicutes pathogens (*Staphylococcus aureus* and *Listeria monocytogenes*), and two species in which limited genetic diversity has previously been detected (*Neisseria gonorrhoeae* and *Mycobacterium tuberculosis*). The pangenome of these datasets were defined using Roary, from which the population-wide core genome was aligned and pairwise distances calculated using the Tamura-Nei (tn93) distance. These pairwise distances use only loci conserved across at least 99% of the isolates in each sample, rather than the pairwise definition of the core intrinsic to PopPUNK. The genome content divergence was measured as the Jaccard distance between the presence and absence of accessory coding sequences. In all cases, there was a strong linear correlation across the full range of both a and π (Fig 3 and Table S1). For π, the linear relationship was close to the identity line, indicating PopPUNK was accurately estimating the per-base probability of sequence divergence using the default sketch size of 10^4^. The exception was *M. tuberculosis*, for which the range of π was an order of magnitude lower than in other species. Therefore PopPUNK needed to be run with a sketch size of 10^5^; despite this increase, the analysis remained fast and memory efficient (Table 2), and the estimated π values strongly correlated with the tn93 distances.

**Figure 3:**
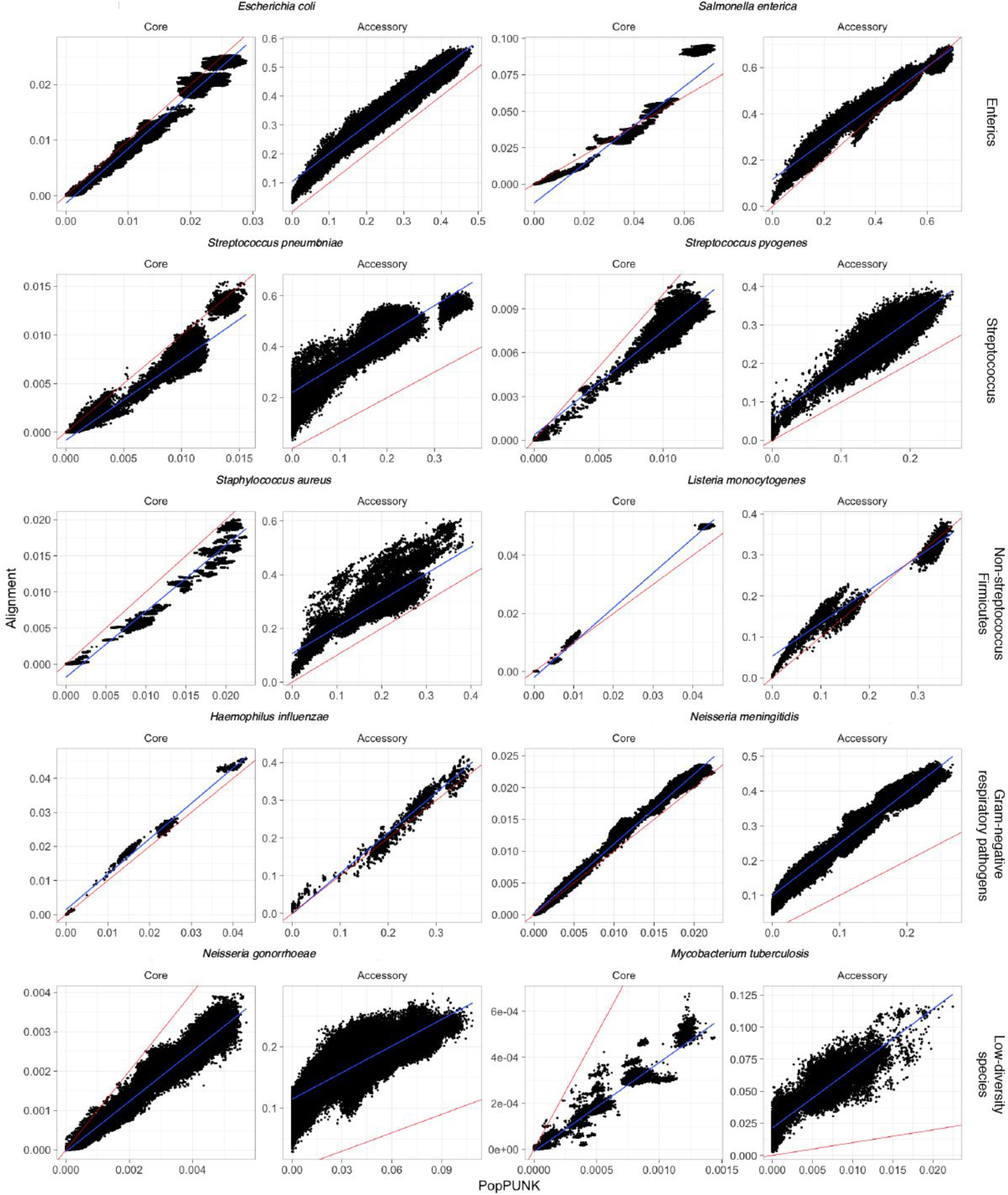
Comparison of core and accessory distances from PopPUNK (x-axis) and pan-genome construction with Roary (y-axis). For each species, the core distance was calculated as the Tamura Nei (tn93) distance from the core genome alignment; the accessory distance was calculated as the Hamming distance between accessory gene presence/absence. In each panel, the line of identity (red line) and a linear regression (blue line) are also plotted.

For a, the best fit line was typically parallel to, but below, the line of identity. The difference in intercept is likely to represent an artefact of the default BLAST identity threshold used by Roary for clustering genes (95%), which causes divergent alleles of orthologous loci to be split into different genes, and the difficulties of automated annotation of draft genome assemblies, which can contain many contig breaks. In the case of S. *pneumoniae*, in which there is a high divergence between the Roary and PopPUNK analyses, the smaller distances calculated by PopPUNK are very similar to those estimated by an independent annotation and analysis of gene content (Fig S4) using COGtriangles (Croucher et al. 2014b; Kristensen et al. 2010). Furthermore, the annotation-independence of PopPUNK means it is not sensitive to variation in the identification of coding sequences that might result from differing assembly quality or inconsistent gene calling.

Manual inspection of genome alignments revealed the discrepancy between the distribution of a in *M. tuberculosis*, inferred to reach up to ~10% by Roary but only ~1.5% by PopPUNK (Fig 3), primarily represents impact of frameshift mutations and the difficulty in consistently assembling and annotating coding sequences within the PE/PPE repeats (Bryant et al. 2013). PopPUNKs ability to align at least parts of these relatively high diversity loci likely account for the higher density of SNPs it infers in the core genome shared by compared strains (Cohen et al. 2015; Bryant et al. 2013). This annotation-independence also means PopPUNK can detect divergence in intergenic regions, increasing its sensitivity to widespread changes in regulatory regions (Oren et al. 2014). Additionally, PopPUNK was between six- and 200-fold faster than Roary, while using between five- and 45-fold less memory. Therefore PopPUNK is an efficient means of accurately measuring SNP and gene content divergence in species-wide genomic datasets. This can also be performed independently of the downstream clustering steps.

### PopPUNK successfully resolves complex bacterial populations into strains

The discontinuous distribution of a and π between all pairs of sequences in a population has previously been shown to reflect the division of the population into strains in S. *pneumoniae* (Croucher et al. 2014b). The distances estimated by PopPUNK successfully replicated this (Fig S3). Such structure within the bacterial population is typically reflected by a separation between the shorter genetic distances, which can be regarded as within-strain comparisons, and the larger between-strain distances, which may form one or more clusters in the plot, depending on the bacterium’s evolutionary history (Fig 2). To test whether this also applied to other bacterial pathogens, the pairwise a and π distributions were plotted for the other nine species-wide collections listed in Table 1 (Fig 4 and S5).

**Table 1:** Adjusted *R*^2^ between PopPUNK inferred core and accessory distances, and core and accessory distances inferred from a core genome alignment produced using Roary. We did not run Roary across the entire *Salmonella* genus. We calculated the Silhouette Index for each sample, using the PopPUNK core and accessory distances and the clustering method indicated in the table header. We show the average Silhouette Index over all samples. The adjusted Rand index representing overlap between PopPUNK clusters and R-hierBAPS clusters is shown, where identical clustering is one and completely different clustering is zero. *For *Listeria monocytogenes* the second level R-hierBAPS clusters are analysed, as the first level represents the deep split between lineages.

| Species | Publication reference | Adjusted $R^2$ | | Number of clusters | | Average Silhouette Index (best = 1, worst = -1) | | Adjusted Rand index |
| --- | --- | --- | --- | --- | --- | --- | --- | --- |
|  |  | Core | Accessory | PopPUNK | R-hierBAPS (level 1/level 2) | PopPUNK | R-hierBAPS (level 1) |  |
| *Staphylococcus aureus*<br>N = 284 | (Aanensen et al. 2016) | 0.96 | 0.69 | 27 | 9/29 | 0.58 | 0.53 | 0.852 |
| *Escherichia coli*<br>N = 1508 | (Kallonen et al. 2017) | 0.98 | 0.97 | 130 | 24/101 | 0.40 | 0.41 | 0.965 |
| *Salmonella enterica*<br>N = 847 | (Alikhan et al. 2018) | 0.91 | 0.97 | 12 | 10/32 | 0.54 | 0.53 | 0.999 |
| * *Listeria monocytogenes*<br>N = 128 | (Kremer et al. 2017) | 1.00 | 0.96 | 31 | 3/18 | 0.60 | 0.55 | 0.924 |
| *Haemophilus influenzae*<br>N = 75 | (Koelman et al. 2017) | 0.98 | 0.95 | 27 | 7/24 | 0.69 | 0.41 | 0.478 |
| *Neisseria meningitidis*<br>N = 882 | (Lees et al. 2017) | 0.99 | 0.96 | 45 | 15/66 | 0.61 | 0.40 | 0.775 |
| *Neisseria gonorrhoeae*<br>N = 1102 | (Grad et al. 2016) | 0.94 | 0.72 | 132 | 15/58 | 0.21 | 0.36 | 0.921 |
| *Streptococcus pyogenes*<br>N = 675 | (Lees et al. 2016) | 0.84 | 0.78 | 167 | 17/61 | 0.76 | 0.18 | 0.102 |
| *Streptococcus pneumoniae*<br>N = 616 | (Croucher et al. 2013, 2015) | 0.83 | 0.75 | 62 | 19/63 | 0.76 | 0.59 | 0.766 |
| *Mycobacterium tuberculosis*<br>N = 219 | (Cohen et al. 2015) | 0.95 | 0.72 | 54 | 7/20 | 0.41 | 0.61 | 0.914 |

**Figure 4:**
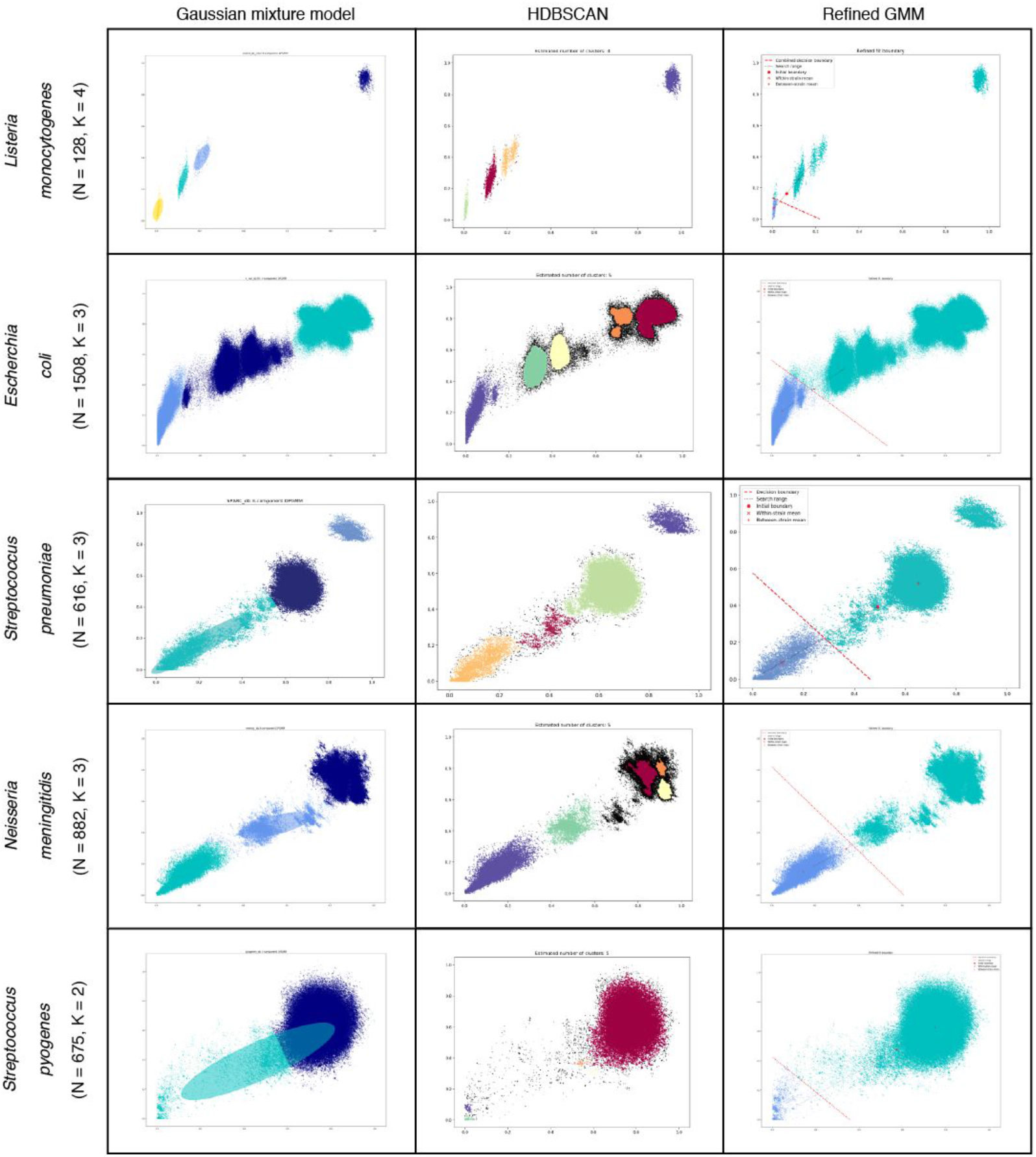
PopPUNK model fitting output for four archetypal examples (other species shown in Fig S5). Each row is a species, with each plot showing the distribution of core and accessory distances, with points coloured by their predicted cluster. The cluster closest to the origin is the within-strain cluster. The two-dimensional Gaussian mixture model (2D GMM) is in the left column, which also shows ellipses with the mean and covariance of the fitted mixture components. The HDBSCAN plot in the centre column additionally shows unclassified noise points as black. The final column shows the fits when maximising the network score to refine the 2D GMM fit. *Listeria monocytogenes* has clearly separated clusters which are well predicted by all methods. Although there is more complex structure on the plots, *Escherichia coii* and *Neisseria meningitidis* have a within-strain cluster also well captured by all approaches. In *Streptococcus pneumoniae* recombination makes the boundary between clusters less distinct, and the mixture model includes too many links (Fig 5). HDBSCAN is more accurate, but the refinement of the initial fit provides the most accurate and intuitive demarcation of the within-strain links. *Streptococcus pyogenes* exhibits low within-strain recombination, hence has a dense cluster of points near the origin of the graph, but high between-strain recombination, resulting in the single, broad between-strain set of points. Network score fit refinement is required for an accurate model fit in this case.

Species known to exchange sequence through homologous recombination at similar frequencies to S. *pneumoniae*, such as *Neisseria meningitidis* (Fig 4), exhibited a similar distribution of pairwise genetic distances. The group of within-strain pairwise distances, found near the origin of the graph, was elongated, likely as a consequence of extensive diversification of strains through transfer of genomic islands and shuffling of core sequence through homologous recombination. The between-strain distances were primarily concentrated within a single dense modal cluster, consistent with the simulations involving high level of recombination in Fig 2. In contrast, *Streptococcus pyogenes* strains exhibit little evidence of recent diversification through homologous recombination (Nasser et al. 2014), hence the within-strain distances were tightly clustered near the origin of the graph (Fig S5). Nevertheless, between-strain distances remained concentrated in a single node, consistent with the much higher level of recombination inferred across broader samples (Didelot and Maiden 2010), resulting in the star-like internal structure of species-wide S. *pyogenes* phylogenies (Chalker et al. 2017). A different pattern was evident in other species in which homologous recombination is infrequently observed, such as *Escherichia coii, Salmonella enterica* and *Staphylococcus aureus* (Fig 4 & S5). This exhibited much stronger evidence of deep population structure, characterised by multimodal between-strain distance distributions that likely reflect ancestral divergences that have not been overwritten by sequence exchange. The distribution of within-strain distances had high variance in the accessory direction, which is likely to reflect rapid diversification in gene content through movement of mobile genetic elements in the absence of core genome diversification through homologous recombination (Zhou et al. 2014; Aanensen et al. 2016; Kallonen et al. 2017). A more extreme version of this pattern was clear in *Listeria monocytogenes* and *Haemophilus influenzae*, which are composed of deep-branching lineages that result in small, tight clusters being formed in the π-a distance space. This structure was apparent even using a small total number of samples. Across a range of ecologically and taxonomically distinct species, there are clear groups separated by both larger and smaller divergences appear on the plot, with the smaller divergences representing within-strain comparisons.

In order to identify these clusters, two alternative approaches are implemented within PopPUNK: two-dimensional Gaussian mixture models (2D GMMs), which split the points into a user-specified maximum of *K* two-dimensional Gaussian distributions, or HDBSCAN, which is run iteratively to identify fewer than a user-specified maximum number of clusters *D* (see Methods). The results of the application of both methods to the genomic datasets listed in Table 1 are shown in Fig 4: in each case, both methods were successfully able to resolve the pairwise distances into discrete clusters. The bacterial population can then be represented as a network in which each node corresponds to an isolate, and each within-strain relationship to an edge between these nodes (Fig 1). This network has the property that strains can be defined as the separate connected components (Fig 5).

**Figure 5:**
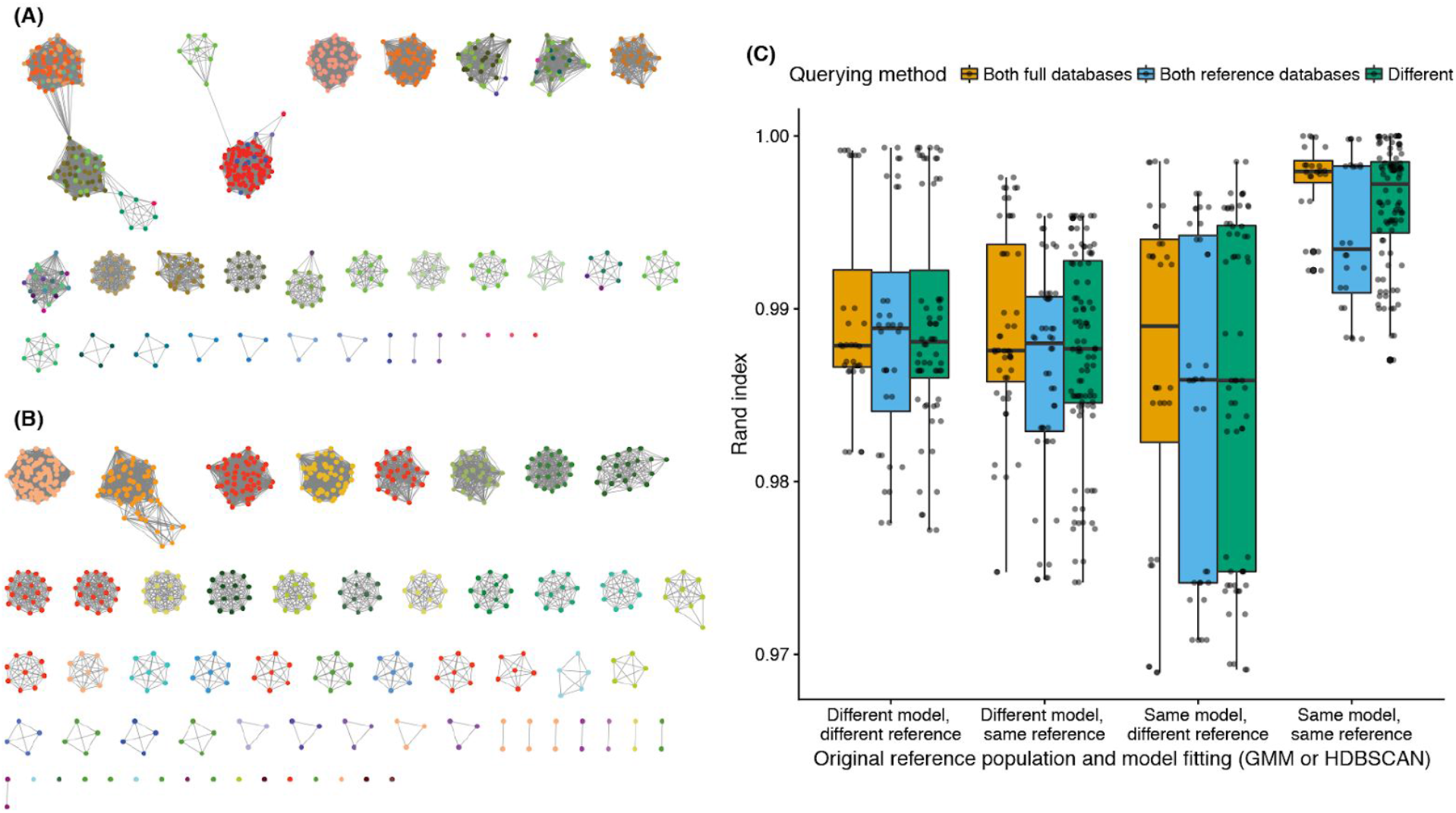
Network and query assignment for S. *pneumoniae*. (A) Cytoscape view of the network for the SPARC dataset using the 2D GMM fit. Nodes (dots) are samples and edges are those pairwise distances classified as ‘within-strain’. The nodes are coloured by clusters for the refined fit in panel (B) showing which clusters are incorrectly merged in the mixture model fit. (B) As in (A), but showing the network after fit refinement. High stress edges causing clusters to be merged have been removed after maximising the network score. (C) Boxplots showing the similarities between cluster assignment when running PopPUNK in different modes. The different model types (2D GMM or HDBSCAN) implemented in PopPUNK were each fitted to either the Massachusetts or Maela S. *pneumoniae* population defined in (Corander et al. 2017), then refined through maximisation of *n*_s_. The three non-reference populations were then added in successive batches, either through comparisons to the full dataset or a representative set of reference sequences selected based on network structure, in all possible permutations. Comparisons of clustering similarity were then calculated between all these permutations in which the final population added was the same using the Rand index (Rand 1971); only those isolates in the most recent extension of the network were used. These values are shown separated according to the starting reference population (Massachusetts or Maela), initial model (2D GMM or HDBSCAN), and comparison method (bar colour; full database or references only).

### PopPUNK can exploit network properties to refine strain definitions

Neither the 2D GMM or HDBSCAN methods alone could satisfactorily resolve the recombinogenic populations into strains, primarily due to the diffuse nature of the within-strain distribution, which likely reflects the heterogenous rates of diversification observed in different strains (Croucher et al. 2013; Didelot et al. 2013). For the 2D GMMs, this was manifested as insufficiently specific, elongated within-strain distributions, which incorrectly included between-strain links as edges. For HDBSCAN, the expectation of a background noise in the distribution meant some pairwise distances between closely-related isolates on the fringes of the main density of within-strain points were omitted from the appropriate cluster. Only a few spurious connections can have a dramatic effect on strain definitions, as a single edge can cause large clusters to be merged, as previously observed for MLST clonal complexes (Turner et al. 2007). Increasing the threshold corresponding to within-strain connections reaches a transition point at which there is a dramatic increase in the density of edges in the network, and a decrease in network transitivity (Fig S6). This represents a small proportion of the many between-strain distances being included as edges, greatly increasing the connectivity of the network, by linking between tightly-knit components with spurious, high-stress edges. However, decreasing the threshold typically reduces the network density while increasing the transitivity of edges (Fig 5A), owing to the strong non-overlapping community structure of the network, meaning components corresponding to strains are highly internally connected. Therefore a network score statistic *n*_s_ ranging between zero and one was defined:

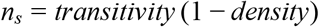

For each dataset, this statistic was first calculated by separating within and between-strain distances. With the 2D GMM this was using a threshold lying on the decision boundary between the within- and between-strain clusters, and with HDBSCAN the point equidistant between the means of the within- and between-strain clusters. The position of this boundary was then optimised to maximise *n*_s_ (see Methods). This provided an intuitive boundary separating the within-strain distances (Fig 4), and tended to be consistent whether initialised from either a 2D GMM or HDBSCAN (Fig S3). Inspection of the network before and after this refinement showed that small numbers of spurious edges between high frequency clusters were removed, and low frequency clusters were kept distinct (Fig 5B). This resulted in a greater number of more robust clusters, and therefore higher clustering specificity.

For the populations listed in Table 1 these strain definitions were evaluated relative to the top level clusters identified from the core genome alignment using R-hierBAPS (Tonkin-Hill et al. 2018). PopPUNK used 15 to 74-fold less memory and ran between ten and 100-fold faster than R-hierBAPS (Table 2). The biggest differences in performance were observed in relatively small collections containing extensive core genome divergence (Fig 3), presumably representing the complexity of fitting a the BAPS model to such data. The number of clusters estimated by each method was similar (Table 1), with an average adjusted Rand index of 0.852 indicating a high level of overlap. Based on the Silhouette distance calculated from the π and a distances, the clustering identified by PopPUNK was of similar, or better, quality than that of BAPS, with the notable exceptions of *N. gonorrhoeae* and *M. tuberculosis*, which lack the assumed strain structure (Fig S4). For instance, in the case of S. *enterica*, the clusters identified agreed perfectly with a recent reappraisal of species and subspecies definitions (Alikhan et al. 2018), whereas using BAPS leads to two cases of subspecies being merged, and a cluster with a single member being added to another larger cluster. This analysis shows PopPUNK is able to efficiently and accurately identify strains within a range of species-wide bacterial population genomic datasets.

**Table 2:**
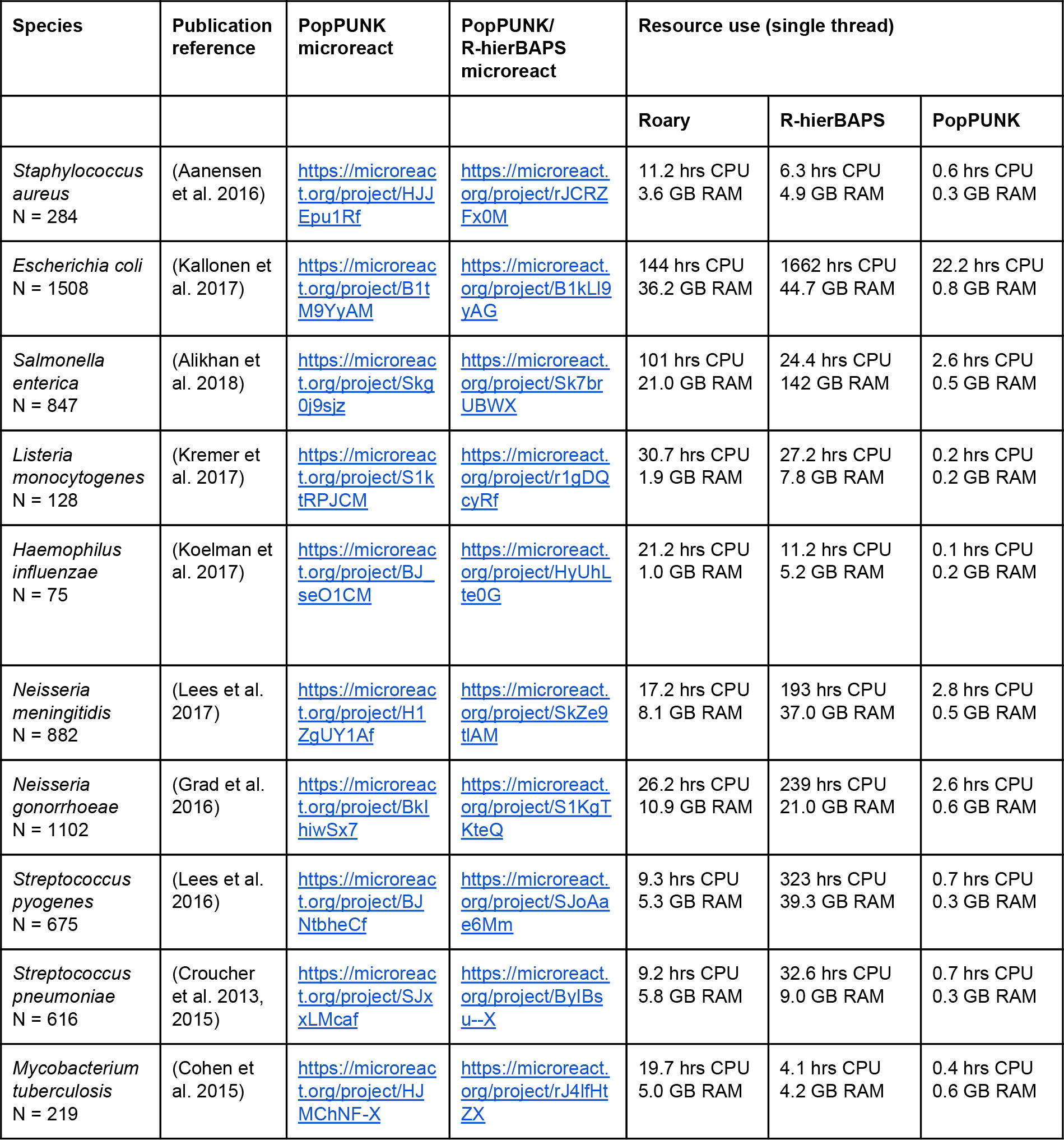
Resource and result comparison for each dataset. For all methods we quote the total CPU time used, and maximum memory. Both PopPUNK and R-hierBAPS use multithreading with close to 100% efficiency, so total wall-time was lower than these estimates roughly by a factor of the number of cores used.

### PopPUNK rapidly integrates new genomic data into clusters

By first generating a reference database and defining a model by which a and π distances can be assigned as being within- or between-strain, PopPUNK allows the network by which strains are defined to be extended. New batches of genomes can then be included without needing to refit the model or recalculate all pairwise distances. This can result in existing strains expanding in number, merging with others, or new strains not previously represented in the database being founded. By automatically rebuilding an updated database, PopPUNK allows iterative expansion through addition of successive batches of genomes. The accuracy of this approach was tested using a dataset of 4,107 draft S. *pneumoniae* genomes resulting from the combination of four different populations, sampled from Massachusetts (USA), Southampton (UK), Nijmegen (Netherlands) and Maela (Thailand) (Corander et al. 2017). Both the 2D GMM and DBSCAN models were fitted and subsequently refined using network properties using either the Massachusetts or Maela collections as the initial reference set. The three non-reference populations were then added as individual batches in every possible permutation to test the consistency of clusters from different starting points. The final clusterings of the isolates in the last population to be added were compared using the Rand index. This metric varies between zero, indicating completely dissimilar clusterings, and one, indicating identical clusterings (Rand 1971). When comparing the different orders in which the non-reference populations were added, using the same reference database and refined models, the Rand indices were all above 0.992 (Fig 5). Hence by expanding the network in this way, PopPUNK can accurately and consistently assign batches of genomes to strains.

Comparisons were also made between queries that were assigned using different initial model fits, to test the sensitivity across the starting parameters of PopPUNK. Whether a different population (Massachusetts or Maela), or a different method (2D GMM or HDBSCAN) was used to initialise the model, there was a slight decrease in the reproducibility of the clustering, although the mean Rand index was still greater than 0.99 (Fig 5). These were all highly similar to the results obtained when all 4,107 isolates were clustered in a single step, regardless of which spatial clustering approach was used (Fig S8). The clustering observed after successive additions of query sequences to the network of genomes therefore exhibits reassuringly little sensitivity to the original choice of population and model fit.

The addition of batches by calculating the distance to every sample in the original clustering is inefficient, as the tight clusters of isolates within the same strain will each be separated from a given query by similar π and a distances, making a high proportion of comparisons redundant. By default, PopPUNK reduces a full database to a set of reference genomes, which includes just one representative from each clique (i.e. a fully-connected component) within the network (Fig 1). This allows for faster and more efficient analysis of new batches of data, from which new references can be extracted through applying the same search for cliques. To test whether this approach caused any decrease in clustering reproducibility, the four test populations of S. *pneumoniae* were successively integrated to the reference Massachusetts or Maela datasets as before, but this time using cliques to pare the database down to a set of references after each expansion of the network. When the cluster assignments of the genomes in the last population to be added were compared between the reference-based approach and with the previous analyses using the full databases, again a high degree of correspondence was measured by the Rand index (Fig 5). Although there was a detectable decline in the consistency of strain assignments using these reference-based searches, this decrease was even smaller than the use of different initial model fits. Correspondingly, only a small decrease in similarity to the single-step clustering was measured (Fig S8). The final networks of all four combined populations contained a median of 256.5 references (range: 212-398): this almost 20-fold reduction in the database size resulted in a median 2.6-fold decrease in CPU time required for the addition of the final batch. Using this network-based design, PopPUNK can thereby efficiently and accurately combine multiple batches of genomes through continually updating a database of representative sequences that span the full diversity of the sampled population.

### PopPUNK outputs can be used directly for interactive browser-based analysis

To facilitate simultaneous analysis of population structure in conjunction with associated epidemiological data, the outputs of PopPUNK can be directly uploaded to the freely-available online epidemiology platform Microreact (Argimón et al. 2016). This is achieved through using the pairwise π matrix to construct a neighbour-joining tree, either natively within PopPUNK using dendropy (Sukumaran and Holder 2010), or by running RapidNJ (Simonsen et al. 2011). The pairwise a matrix is used to generate a t-SNE projection using the sklearn package (Pedregosa et al. 2011), which is displayed using Microreac?s network interface developed for PANINI (Abudahab et al. 2017). Links to examples of such analyses are provided in Table 2. We have also included options for outputting the results of PopPUNK in formats appropriate for the online visualisation software Phandango (Hadfield et al. 2017) and GrapeTree, the browser-based viewer used in Enterobase (Zhou et al. 2017). The underlying network can also be investigated using Cytoscape (Shannon et al. 2003).

The combined results arising from querying one dataset against another using PopPUNK can also be displayed with these platforms, with the additional isolates highlighted in the output. This can be used for iteratively merging similarly-sized datasets, as illustrated in Fig 5C (an example of such a merging in S. *pneumoniae*: https://microreact.orα/oroiect/SkZ23iPbX). This means PopPUNK can be applied as a rapid tool for ruling out potential outbreaks. As an example, we queried 175 S. *pneumoniae* isolates from the multidrug-resistant PMEN14 lineage (Croucher et al. 2014a) against the diverse Massachusetts species-wide carriage population sample of 616 isolates (https://microreact.org/oroiect/BkNqKdPb7). This identified all the query isolates as belonging to a single strain, and generated a phylogeny in which they were confined to one clade and an accessory projection representing gene content differences, in less than six minutes using sixteen CPUs and less than 200 MB RAM. Repeating this analysis using an optimised reference database of 63 representative sequences, the same process completed in under four minutes (https://microreact.org/oroiect/Hk-_F0oWX). Hence PopPUNK provides a simple and efficient means to intuitively and interactively analyse complex data using a platform that facilitates online collaboration and publication.

### PopPUNK can be used for analysis of low diversity pathogens

In the cases of *N. gonorrhoeae* and *M. tuberculosis*, R-hierBAPS generated substantially better clustering than PopPUNK, as judged by the Silhouette index. This likely represents the absence of true strains in these species, which are homogeneous compared with the other species studied, and therefore do not exhibit the discontinuous but correlated divergence in π and a assumed in the clustering stage. In *N. gonorrhoeae*, the accessory genome content consists of a small number of prophage, three plasmids and the 80 kb gonococcal genomic island (GGI), all of which can vary independently of core genome divergence (Morse et al. 1986; Hamilton et al. 2005; Bennett et al. 2010). Using the fit based on the network score, we were able to successfully split the population into 132 clusters. Inspection in Microreact revealed a clade composed of a polyphyletic mixture of clusters 5 and 10. Their close relationship within the core genome tree suggested the difference in clustering reflected divergence in their accessory genome. To identify the specific loci responsible for this split, microbial genome-wide association software (pyseer) was used with these isolates, finding 3,679 *k*-mers distinguishing the clusters 5 and 10 (Lees et al. 2016, 2018a). The top hits recovered following mapping of these *k*-mers to reference sequences were the GGI and phage sequence, confirmed as being the distinctive loci through manual inspection.

**Figure 6:**
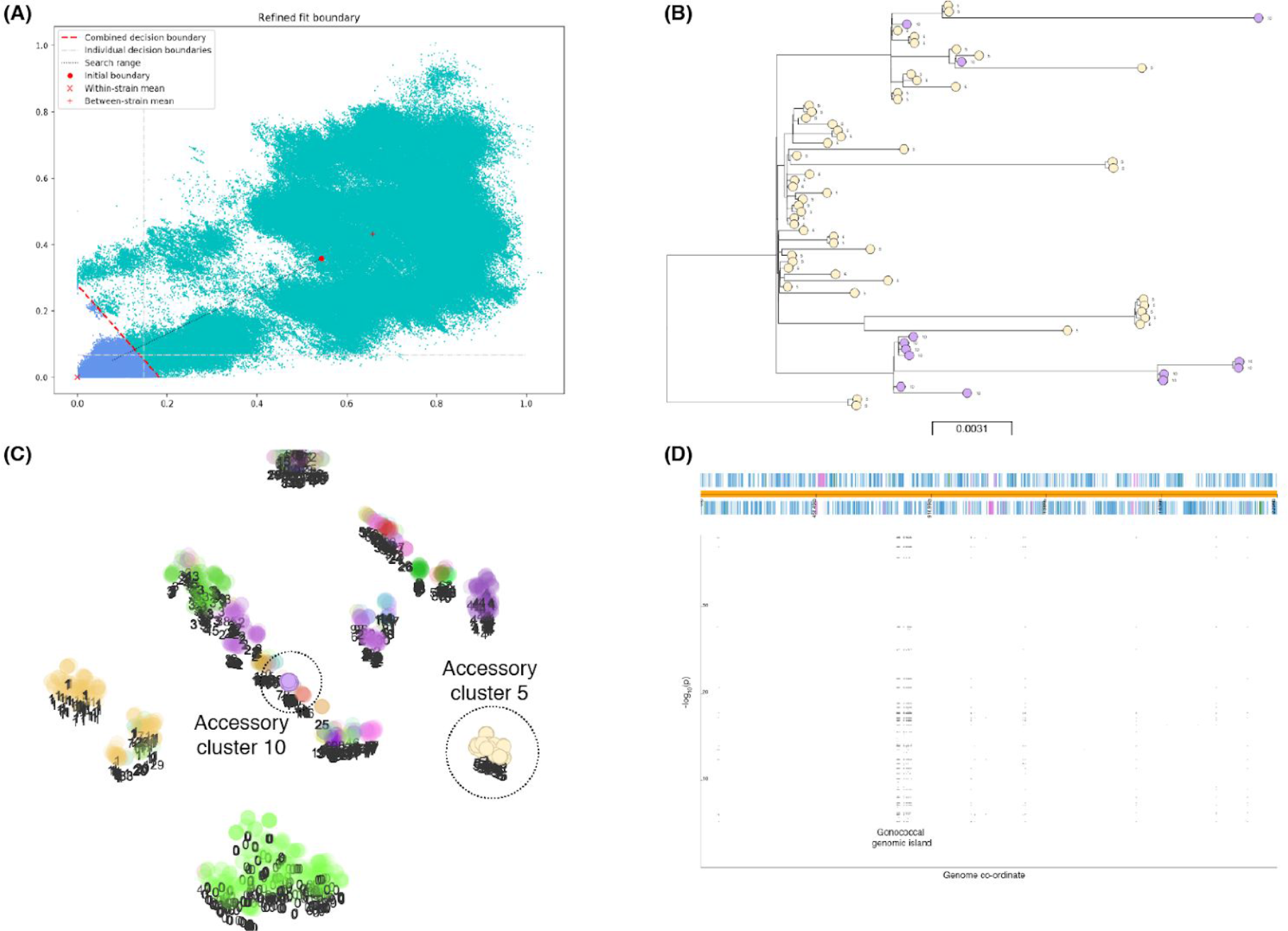
PopPUNK analysis of *Neisseria gonorrhoeae*. (A) Result of optimising the position of the decision boundary on the combined distances (red dashed line), as well as core and accessory only (dashed gray lines). These methods resulted in 132, 114 and 92 clusters respectively. (B) Example of a single core cluster, which is split into separate clusters using either the combined or accessory boundary. Shown is the phylogeny, with tips coloured by combined cluster. (C) Accessory t-SNE, with the clusters in b highlighted, confirming they diverge. (D) Manhattan plot of *k*-mers associated with the subcluster split in panel (B), which show the GGI is responsible for this difference.

For cases where there is independent core and accessory evolution, such as within a lineage, we therefore also implemented a more suitable model which just uses one of the core or accessory distances, rather than a combined score. All three clusterings can be jointly inspected using the Microreact output (for example https://microreact.org/oroiect/S1KgTKteQ). In this particular cluster, the core distances no longer separated the isolates with the GGI and prophage, whereas the accessory distances gave a similar clustering to the combined boundary.

Applying the same analysis to *M. tuberculosis* (https://microreact.org/oroiect/rJ4lfHtZX) demonstrated the PopPUNK phylogeny accurately reconstructed the previously-identified lineages in this population (Cohen et al. 2015). While the top-level R-hierBAPS effectively identified these lineages, the PopPUNK core genome sequence clusters were much more finely grained, resembling the categorisation into spoligotypes, which are informative for more detailed epidemiology. As PopPUNKs clusters can be easily assigned to lineages or R-hierBAPS clusters using the core phylogeny, this ensures the high-resolution links identified using this software can also be used to for analysis at broader levels.

Such detailed epidemiology can also be valuable within more diverse species, such as when analysing of individual strains. As an example of such an hierarchical study of within-strain diversity, a PopPUNK analysis of the S. *pneumoniae* PMEN14 lineage (Croucher et al. 2014a) was compared to previous accessory genome study using PANINI, which applies t-SNE to the accessory gene presence/absence matrix generated by Roary (Abudahab et al. 2017) (Fig S9). Due to the lower sequence divergence within a single lineage, PopPUNK was run with sketch sizes of 10^4^ (https://microreact.org/proiect/H1UsF5CxX) and 10^5^ (https://microreact.org/project/H1Av59CIQ). In both cases, the phylogeny and accessory clusterings were similar, with the t-SNE projection recapitulating the main results from the PANINI analysis: group 5, lacking the Tn*916* antibiotic resistance element, were resolved as being separate from the rest of the population, while groups two and three, both carrying two prophage, were separated from the non-lysogenic group one, despite them being polyphyletic in the core genome tree.

### Discussion

Current typing and model-based clustering methods do not fulfil all the needs of a population analysis tool for bacterial genomic epidemiology. Here we have introduced PopPUNK, a novel suite of algorithms for analysing large species-wide bacterial population genomic datasets overcoming the technical and computational limitations of previous approaches. Our methods are based upon the rapid estimation of pairwise distances between isolates both in terms of divergence between their shared sequences and differences in their gene content. The use of MinHash optimised *k*-mer comparisons, which are independent of annotation and alignment, means these methods are highly efficient in both memory and CPU usage. Compared to construction of a core-genome alignment, our method is up to 200 times faster and produces comparable results. The use of *k*-mer comparisons also fully exploits the information available in the entire genome assembly, without being limited to comparisons with a predefined set of common loci. These distances provide a clear overview of the population structure and the evolutionary mechanisms through which it is shaped, reflecting the degree to which the population is split into deep lineages, the scale of accessory genome variation between isolates, and the inferred effects of recombination on population structure. By altering the MinHash sketch sizes, PopPUNK users can optimise for speed, with default parameters rapidly identifying strains in diverse species, or precision, up to the level of single nucleotide resolution in the pan-genome. The software is also flexible in its application of machine learning techniques to these distributions of distances, making it applicable across a wide variety of population structures and collection sizes. As these spatial-clustering approaches are refined, and new methods become available, PopPUNKs modularity means the repertoire of these techniques used by the software can be further expanded.

For species with discontinuous π-a distributions, categorisation of the population into clusters of related isolates is epidemiologically critical both for following longitudinal trends and understanding the distribution of clinically-important traits in cross-sectional samples. Depending on the nature of the pathogen, PopPUNK can divide collections into sequence clusters, using the divergence between shared sequences (Palys et al. 1997); genomotypes, defined as being similar in the accessory loci they harbour (Doolittle 2002); or strains, here defined as those isolates similar in both their core genome and gene content. The consistency of PopPUNKs clusters with those identified by BAPS, which in turn match well with those identified by MLST and wgMLST (Alikhan et al. 2018), emphasises the likelihood that such clusters represent coherent natural populations. In each mode, PopPUNK applies a stringent threshold that minimises the probability of spurious links, corresponding to edges with high betweenness, which eventually lead to ‘straggly’ clusters containing distantly-related isolates linked indirectly through intermediates. This is achieved both through careful identification of the within-strain distances, and pruning the resulting networks such that the proportion of transitive relationships is maximised. This approach also avoids the problem of clusters arising from diverse sets of rare genotypes, rather than lineages descended from a recent common ancestor (Grad et al. 2016; Willemse et al. 2016; Croucher et al. 2013). PopPUNK instead separates these into multiple small, distinct groupings, emulating one of the most desirable properties of a high quality MLST scheme.

Importantly, the stringency of the within-strain threshold also ensures strain definitions are persistent, and therefore robust to the addition of further batches of data, as demonstrated by the analysis of different combinations of four globally distributed S. *pneumoniae* datasets. PopPUNK is therefore ideally suited to addressing the current limitations of *k-*mer-based epidemiological methods which suffer from the absence of an appropriate curated database or a stable strain nomenclature (Nadon et al. 2017). The outputs can be readily shared using the browser-based Microreact platform, and PopPUNK databases can be downloaded, queried and extended by different users in a non-centralised manner, allowing new batches of data to be continuously integrated in ongoing analyses. The speed of our software makes it capable of handling the thousands of bacterial genomes that will routinely be generated through surveillance of common bacterial infections. This is a consequence of the fast annotation- and alignment-free sequence analysis methods used, and the integration of data into a network structure, which allows the overall population to be represented by a small number of reference sequences. Our software has the potential to underpin global online surveillance databases for bacterial pathogens, which is particularly relevant as public health agencies around the world seek to fully exploit the benefits of genomic epidemiology.

## Methods

### Rapid calculation of core and accessory divergences

PopPUNK uses pairwise comparisons through *k*-mer matching between two sequences (s., and *s*_2_) at multiple *k* lengths to distinguish divergence in shared sequences, π (Nei and Li 1979), from divergence in accessory locus content, here defined as a, the Jaccard distance between the sequence content:

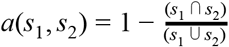

For any given *k*, a MinHash algorithm can efficiently estimate a,. However albeit with confounding by divergence due to π that prevents matching between *k*-mers in the core genome common to (*s*_1_,*s*_2_) is confounding. Assuming that such sequence mismatches are distributed evenly throughout the genome, a can be estimated independently of *k* and π by calculating a function for each (*s*_1_,*s*_2_) pair that relates the proportion of shared *k*-mers *p*_match_, π, and a over a series of *k*-mer lengths *k*:

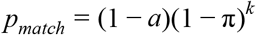

Which we fit as a linear relationship in log space by minimizing the least squares divergence, and constraining a > 0; π > 0:

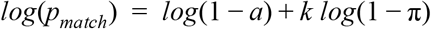

As both distance estimates are symmetrical, only a single comparison is calculated between each (*s*_1_, *s*_2_) pair, corresponding to the upper triangle of a square distance matrix, or (*n*-1)…*n*/2 comparisons. We use Mash (Ondov et al. 2016) with a default sketch size of 10^4^ to efficiently calculate *p*_match_ for every second *k*-mer size from *k* = *k*_min_ to *k* = *k*_max_ (29, by default). We choose the minimum *k*-mer size *k_min_* for longer sequences such that the probability of a random *k*-mer match *p*_random_ is less than 5%:

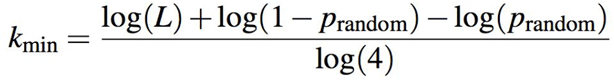

Where *L* is the genome length and log(4) enters due to the alphabet size (assuming minimal gaps or unspecified bases). For typical bacterial genomes with *L* between 1-8 Mb, this corresponds to a *k_min_* of either 12 or 13.

### Automated classification of within-cluster distances

We implemented two models to classify which distance pairs (π, a) are within the same cluster, the choice of which depends on the dataset being used. The first fits a two-dimensional Gaussian mixture model (2D GMM) to a random subsample of up to 10^5^ distance pairs using variational Bayes inference. We use the default implementation in scikit-learn 0.19 with a Dirichlet Process prior on weights, choosing the best final likelihood from five *k*-means initial starts. The maximum allowed number of mixture components (K) can be specified by the user, depending on the plotted distribution of pairwise π and a distances; by default, *K* is set to two. We use membership of mixture component closest to the origin (checking that it contains at least one point in the reference database, as the Dirichlet process prior allows mixture component weights to be set ≃ 0) to define within-cluster distances. All distance pairs are then classified with the fitted model.

The alternative approach uses HDBSCAN to classify a subsample of 10^5^ points using the Boruvka ball tree algorithm (Mclnnes et al. 2017). This set of points is iteratively analysed with progressive reductions in both the minimum number of samples, which defines the how conservative the clustering is, and the minimum cluster size, which determines the threshold number of points a cluster must contain, until there are fewer than *D* clusters (100, by default), and extent of the points in the cluster closest to the origin (assumed to represent within-strain distances) do not overlap with those in the most numerous cluster (assumed to represent between-strain distances) on either axis.

### Use of within-cluster distances to define a reference network and clusters

We use networkx v2.1 (Hagberg et al. 2008) to construct an undirected graph with unweighted edges to define population clusters. Each sample in the reference database is a node in this graph, and distances classified as within-cluster by the above model are added as edges between the corresponding nodes. Clusters are then simply defined by extracting the connected components of this network, and ordered by the number of isolates they contain, from largest to smallest. Evaluation of the network structure uses a score, *n*_s_:

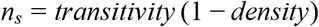

This score ranges between zero and one, with values close to the upper bound corresponding to a good fit, as every isolate in a cluster should share a within-cluster link to every other member of the cluster. This is achieved in spite of the sparseness enforced by the (1 - density) term, which is necessary to subdivide the overall population (Fig S6). After definition of clusters, we randomly select just one member of each clique in the network (sets of nodes where each member node is mutually connected to every other member node) to use in an updated reference database. This removes redundancy in the distances that need to be calculated for database querying, and speeds the assignment of further batches of data.

### Refinement of distance classification using network properties

Both the 2D GMM and HDBSCAN rapidly and robustly identified the main clusters of within- and between-strain distances in π-a space, which were typically well-resolved. However, being conservative in assigning points to the within-strain cluster is not intrinsic to these methods, which treat clusters symmetrically. The relatively small numbers of false positive within-strain assignments from the low density of points between clusters strongly impacted upon network structure by linking components, and consequently the strain definitions. Therefore a model refinement mode was developed to precisely delimit the range of π-a distances that were treated as within-strain links, in order to maximise *n*_s_. If a 2D GMM or HDBSCAN model has already been fitted, then we construct a line between the means of the within- and between-strain clusters, then draw a decision boundary normal to this line (Fig 1). If neither model fitted satisfactorily, then the mean positions of the within- and between-strain cluster means can be provided manually.

We then allow this triangular boundary, distinguishing within- and between-strain distances, to be moved over a user-set range forward and backward from the starting point. We globally maximise *n*_s_ first by testing the network score when placing the boundary at 40 equally spaced points over the allowed range. Local maximisation near this global optimum is then performed using Brent’s method. We have also made it possible to run this optimisation with a vertical boundary (using core distances, π, only) or a horizontal boundary (using accessory distances, a, only) if desired.

### Defining the cluster of query sequences using a previous reference database

New sequences can be rapidly assigned to either a pre-existing cluster, or start to form a new cluster, by addition to the reference network. First, distances are calculated between each query and each member of the reference database using the variable *k*-mer method as above. New queries are added as nodes in the network, and those distance pairs classified as within-cluster are added as edges. As before we define clusters as the connected components of this network, while ensuring labelling consistency with the reference fit. By only calculating reference-query distances rather than all distances, we can perform assignment of *M* queries using a reference database with *N* members using *NM* distance calculations rather than (*N*+*M*)^2^. If the database is being updated for further querying, all *M*^2^ query-query distances are also calculated (for a total of *NM* + *M*^2^ distances) so that cliques in the network can still be used to extract representatives for each cluster. However, in practise the construction of *M* sketches is the most computationally expensive step, which gives roughly linear query time in both cases.

### Output and visualisation

We automatically produce plots to diagnose dataset characteristics, quality of model fit and assignment of distances to clusters using matplotlib v2.1.2 (Hunter 2007). For the distances selected for model fitting, we plot contours of a kernel density estimate using an Epanechnikov kernel. This can also be used to identify outliers with contamination; we provide a program to remove these isolates from reference databases. Plots are generated for each model type showing details of its fitted parameters, together with cluster assignment of the distances. For 2D GMMs, we also plot equal likelihood contours and the decision boundary for within- and between-cluster assignments (Fig S3).

While a neighbour joining tree constructed using pairwise Jaccard distances directly has been shown to be reasonably accurate, using core genome divergence gives a more accurate tree topology (Lees et al. 2018b). We therefore use dendropy or RapidNJ to produce a midpoint rooted neighbour joining tree from the core distances (Simonsen et al. 2011; Sukumaran and Holder 2010), and t-SNE to perform clustering of accessory distances (Abudahab et al. 2017). To enable interactive visualisation of these outputs, PopPUNK can write files formatted for Microreact (Argimón et al. 2016), Phandango (Hadfield et al. 2017) and GrapeTree (Zhou et al. 2017). Each of these can be automatically joined with other user-provided metadata for visualisation. We also produce output for Cytoscape (Shannon et al. 2003) for inspection and analysis of the network.

### Code optimisation for large datasets

We optimised our code such that datasets with up to ~10^4^ samples could be analysed in a single step. Where possible, we used numba vθ.36.2 to compile functions (Lam et al. 2015) and exploited multithreading of Mash sketching and distance calculations. We also multithreaded the regressions to calculate core and accessory distances, using the sharedmem package (vθ.3.5) to avoid copying and storing large distance matrices in main memory (Feng et al. 2017). We infer sequence labels of rows in distance matrices by their order, rather than storing them in memory. For larger datasets, fitting a reference model to a subset of samples, then adding in query sequences iteratively makes analysis tractable.

### Comparison with other methods using both simulated and real data

To determine the specificity of PopPUNK in distinguishing sequence divergence from differences in gene content, forward-time simulations were run using BacMeta (Sipola et al. 2018). A population of 1000 bacteria, each represented by 100 loci each one kilobase long, was simulated for 1000 generations. Insertions and deletions were fixed at a length of 250 bp. Recombinations always exchanged a complete locus, and were independent of sequence divergence between donor and recipient. A sample of 25 genomes were output from the final generation of each simulation, which were analysed using PopPUNK using default settings. Pairwise distance estimates from 50 independent simulations were then combined for plotting.

To compare with other clustering methods, we selected a range of previously published datasets on ten different bacterial species (Aanensen et al. 2016; Kallonen et al. 2017; Kremer et al. 2017; Koelman et al. 2017; Lees et al. 2016; Grad et al. 2016; Lees et al. 2017; Croucher et al. 2013, 2015; Alikhan et al. 2018; Cohen et al. 2015). For each dataset, as well as PopPUNK, we ran Roary (Page et al. 2015) to construct a pan-genome, using a BLAST sequence identity cutoff of 95%. We calculated core distances using the Tamura Nei (tn93) distances in the core genome alignment. For accessory distance, we used the Jaccard distance between the accessory gene presence/absence vectors. For comparison with another high-performance clustering algorithm, we ran the R version of hierBAPS (https://github.com/gtonkinhill/rhierbaps) using between 8-16 cores depending on dataset size. We estimated the maximum cluster size by dataset, using the output from Roary and information from published analyses of these datasets. For qualitative comparison in Microreact, for each species we also generated a maximum-likelihood tree from the core genome alignment SNPs using IQ-TREE v1.6 with a GTR+I+G+ASC model (Nguyen et al. 2015).

For species with a monomorphic population structure there is not necessarily a clear correlation between a and π. In this case it is logical to define independent sets of clusters using the two distances separately. An example of this is *Neisseria gonorrhoeae*, which we describe fully in the Results. We used a subset of isolates, all of which were in the same cluster based on π but contained two different clusters based on a. We performed a genome-wide association study to determine the specific genes responsible for the two different a-based clusters using pyseer v1.1.0 to associate *k*-mers counted from the entire π-based cluster, and using the a-based cluster as the phenotype. Default settings were used, with the kinship matrix generated using the maximum-likelihood tree under a mixed effect association model (Lees et al. 2018a). We mapped the significant *k*-mers to two reference genomes, which between them contain all known accessory elements for *N. gonorrhoeae*.

### Data access

Code is available on github https://github.com/johnlees/PooPUNK (Apache 2.0 license) and through the python package index (PyPI). Documentation can be found on readthedocs http://poppunk.readthedocs.io/. Online and interactive Microreact instances produced for each data set are listed in table S1. PopPUNK databases with the best model fits for each species can be downloaded from https://doi.ora/10.6084/m9.fiashare.6683624.

## Acknowledgements

We thank Nabil-Fareed Alikhan for helpful discussions about the *Salmonella enterica* dataset, and Leon Bentley for early experiments with changing mash *k*-mer and sketch sizes.

## Author contributions

Conceptualization: JAL, NJC. Data curation: JAL, SRH, RAG, SL, NJC. Formal analysis: JAL, NJC. Funding acquisition: JNW, SDB, NJC. Investigation: JAL, NJC. Methodology: JAL, NJC. Software: JAL, GTH, JC, NJC. Supervision: JNW, JC, SDB, NJC. Validation: SRH. Visualization: JAL, NJC. Writing – original draft: JAL, NJC. Writing – review & editing: All authors.

